# A graph-matching based metric of functional connectome distance between pairs of individuals varies with their ages, cognitive performances and familial relationships

**DOI:** 10.1101/2022.10.03.510660

**Authors:** Hussain Bukhari, Chang Su, Elvisha Dhamala, Zijin Gu, Keith Jamison, Amy Kuceyeski

## Abstract

Functional connectomes (FCs), represented by networks or graphs that summarize coactivation patterns between pairs of brain regions, have been related at a population level to age, sex, cognitive/behavioral scores, life experience, genetics and disease/disorders. However, quantifying FC differences between pairs of individuals also provides a rich source of information with which to map to differences in those individuals’ biology, experience, genetics or behavior. In this study, graph matching is used to create a novel inter-individual FC metric, called swap distance, that quantifies the distance between pairs of individuals’ FCs. We apply graph matching to align FCs between pairs of individuals from the the Human Connectome Project (N = 997) and find that swap distance i) increases with increasing familial distance, ii) increases with subjects’ ages, iii) is smaller for pairs of females compared to pairs of males, and iv) is larger for females with lower cognitive scores compared to females with larger cognitive scores. Regions that contributed most to individuals’ swap distances were in higher-order networks, i.e. default-mode and fronto-parietal, that underlie executive function and memory. These higher-order networks’ regions also had swap frequencies that varied monotonically with familial relatedness of the individuals in question. We posit that the proposed graph matching technique provides a novel way to study inter-subject differences in FC and enables quantification of how FC may vary with age, relatedness, sex and behavior.

## 1 Introduction

Since the discovery of blood oxygenation level-dependent functional MR-imaging (fMRI) around 30 years ago, much of human neuroimaging research has focused on quantifying and analyzing *in vivo* brain activity patterns in humans. Recent analyses of large-scale fMRI datasets like the Human Connectome Project (HCP) have revealed that the functional connectome (FC), or temporal coactivation patterns between different brain regions, is likely heritable, modified by experience, and related to anatomical white-matter connectivity, age, sex, behavior/mood, breathing patterns, wakefulness and physical and cognitive functioning^1–18^. Many of these discoveries rely on brain-behavior mapping approaches that use statistical group-level testing or regression models to relate FC metrics and individual traits across large cohorts of subjects^19–22^. However, population-level group comparisons and correlations sacrifice a rich source of information about inter-individual variability - differences between pairs of individuals. Further, brain-behavior mapping studies of FC many times use atlas-based approaches that depend on a single brain parcellation, derived from anatomical or group-level functional activation information, that is identical for (or very similar across) each individual. It is well known that individual brain anatomy and physiology can vary widely across people - thus, individual-based brain parcellation schemes were developed that considered a person’s unique anatomy and physiology^23–25^. These types of approaches can be difficult to use in brain-behavior mapping, however, as between-subject correspondence of brain regions is no longer straightforward.

A few recent works have shed light on quantifying individual-level differences in FC matrices. For example, it was shown that an individual’s FC is identifiable across sessions, primarily due to individual-level fingerprints in higher order systems like the fronto-parietal network (FPN)^26–28^. This stability of neural dynamics within the same individual has been confirmed across different scan sessions months to years apart, and across different scanners/ sites^14,25,29,30^. Moreover, FC patterns exhibit higher correlation between siblings than unrelated pairs of individuals - an effect which is driven by brain regions again belonging to high order regions like the FPN and default mode network (DMN)^11,31^. Other works have compared individual FCs to a group-average FC, or pairs or triplets of individuals’ FCs, with the goal of classifying individuals into groups defined by specific psychological traits or brain disorders^32,33^. Some studies have performed temporal realignment of regional fMRI time series across subjects to understand the relationship between inter-subject differences in metrics like FC density by subtracting individual values from a group-based average^34^. In subsequent analysis, the difference between the synchronized time-series and real time-series of a subject as a pair-wise metric of FC is used as an input for regressions that ascertain group differences across disease states^35^. Yet, studies conducted on the individual or inter-subject level on a large population, or those that seek to study the variability of FC patterns between two subjects are relatively uncommon^36,37^. To our knowledge, we are the first to focus on how the pair-wise differences in FC among a large, healthy population of individuals maps to the pair of individuals’ respective ages, cognitive performances and familial relationships.

The pair-wise FC metric we employ here is based on an algorithm called graph matching. Graph matching is a mathematical process wherein a permutation matrix is identified that, when applied to a given graph or network, maximizes the correlation between that graph and another target graph^38,39^. Recent work from Osmanlioğlu and colleagues used graph matching to assess the alignment of white matter structural connectivity (SC) and FC across healthy individuals, and found 1) more alignment in somato-motor and visual areas and less alignment in higher-order areas and 2) less alignment of structure and function across development^40^. For our recent work^41^, we applied graph matching to precision-based FC matrices to track longitudinal functional reorganization following stroke. There, we found that regions with more SC damage due to the stroke had more functional reorganization (measured using graph matching), and, further, that individuals having more functional reorganization over time also had larger gains in motor function. These studies point to the utility of graph matching applied to FCs to reveal behaviorally and anatomically relevant information, however, this approach has not yet been applied to investigate inter-subject FCs across a large population of individuals.

In this study, we use graph matching to understand the variability between pairs of individuals’ FCs across young adulthood using data from the Human Connectome Project’s Young Adult (HCP-YA) dataset^3^. Here, our graph-matching technique uses an atlas-based FC from one individual and personalizes it via permutation to maximize correspondence with a second individual’s atlas-based FC; this procedure enables quantification of FC differences between pairs of individuals^39,41–43^. We then relate this individual-pair FC measure to the pairs of individuals’ familial relationship, ages, sexes and cognitive scores. Finally, we quantify how each region or network contributes to the graph-matching distance metric, revealing which networks or regions vary the most across pairs of individuals.

## 2 Methods

### 2.1 Subjects, MRI acquisition, preprocessing and functional connectome construction

We used the minimally preprocessed resting-state fMRI data^3^ from 997 individuals (22-37 years old, 532 female) from the 1200 subjects Human Connectome Project (HCP) data release; these 997 subjects were those that had complete MRI data (including 4 complete fMRIs) that passed quality control. The final set of subjects contained 130 pairs of monozygotic twins, 95 pairs of dizygotic twins and 200 pairs of full siblings. A subset of 41 subjects (out of the 997) had test-retest data consisting of two MRI scans ~6 months apart. Briefly, MRI data was acquired via a Siemens Skyra 3T scanner and included T1- and T2-weighted anatomical MRI (0.7mm isotropic) and resting-state fMRI (2.0 mm isotropic, TR/TE = 720/33.1 ms, 8x multiband acceleration, 14 minutes per scan, 4 scans total over two days)^3^. The minimal pre-processing pipeline performed by the HCP consortium included motion and distortion correction, registration to subject anatomy and standard MNI space, and automated removal of noise artefacts by independent components analysis^44–46^. We parcellated the brain using the Craddock 400 (CC400) atlas^23^, which used spatially constrained spectral clustering applied to fMRI data to partition the brain into 392 regions. Each region was assigned to a Yeo functional network delineating visual, somatomotor, dorsal attention, ventral attention, limbic, default mode, and fronto-parietal networks^47^; we also added a subcortical and cerebellar network for whole brain coverage as in previous work^48^.

We regressed the global signal and its temporal derivative from each subject’s time series, and derived the zero lag Pearson correlation between the four scans’ concatenated time series from each pair of regions. Precision-based functional connectivity (FC) matrices were then obtained by calculating the Tikhonov-regularized inverse of these Pearson correlation matrices^18,49^. Precision FC matrices were used as they have the advantage of being more sparse and reflective of direct connections, which may reduce inter-subject variability. To identify the optimal regularization parameter for the matrix inversion, first an unregularized group-level precision matrix was constructed by inverting the group-average correlation matrix without regularization. The regularization term was set at the value which minimized the sum of the Frobenius norm between each individual’s regularized precision matrix and the unregularized group precision matrix (identified via a grid search over 100 candidates in the log space from 0 to 1).

### 2.2 Graph matching procedure

Let **A**_*i*_, **A**_*j*_ ∈ ℝ^*p*×*p*^ (*p* = 392) denote precision FCs for individuals *i* and *j* computed as described in Section 2.1. The goal of graph matching is to identify a permutation matrix Π ∈ {0,1}^*p*×*p*^ such that the Frobenius norm between the permuted matrix Π^⊤^**A**_*i*_Π and **A**_*j*_ is minimized. Equivalently, this permutation matrix Π, when applied to the vertices of a given graph or network, maximizes the similarity between that graph and a target graph. Here, Π is a binary matrix with exactly one non-zero entry in every row and every column, and Π_*kl*_ = 1, *k* ≠ *l* indicates a permutation (or swap) from region *k* of **A**_*i*_ to region *l* of **A**_*j*_.

Precision FC matrices contain noise from many sources, including noise in the fMRI acquisition and noise in the precision matrix calculation. Direct minimization of an unregularized discrepancy measure between Π^⊤^ **A**_*i*_Π and **A**_*j*_ can lead to a large number of swaps in Π due to this noise in the FC rather than true inter-subject variation, rendering the metrics that we subsequently develop less informative. In particular, we observed a disproportionate number of swaps within and between the limbic, subcortical, and cerebellar networks (areas that are known to have lower signal-to-noise ratios in fMRI) when matching the same individual’s test-retest FCs (Supplemental Figure S1A). To combat this issue, we regularized the estimate of Π. First, motivated by the observation regarding the larger number of swaps in the limbic, cerebellar and subcortical networks in the test-retest data (compared to regions in other functional networks), we introduced two possible penalties to be incorporated into the graph matching optimization procedure. The first (more strict) penalized swaps that involved a limbic, subcortical and cerebellar region, regardless of the other region’s network assignment. This penalty was encoded in the matrix Ω ∈ ℝ^*p*×*p*^ where **Ω**_*k,l*_ = 1 if *k* ≠ 1 and region *k* or region *l* belonged to any of the three networks, **Ω**_*k,l*_ = 0 otherwise. The second (less strict) penalized swaps between region-pairs where both regions belonged to either limbic, subcortical or cerebellar networks. This penalty matrix, Ω ∈ ℝ^*p*×*p*^ had **Ω**_*kl*_ = 1 for pairs of regions (*k, l*), *k* ≠ *l* where both regions *k* and *l* were in limbic, subcortical and cerebellar networks. See see Supplemental Figure **??**B for a visualization of these two penalty matrices. We then minimized the regularized graph-matching objective function

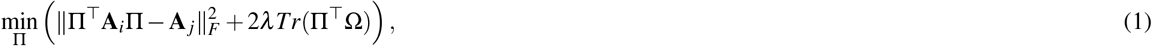

where *Tr*(·) denotes the trace, || · ||_*F*_ denotes Frobenius norm, *λ* ∈ ℝ is the penalty weight, and Π ∈ {0,1}^*p*×*p*^ is constrained to be within the set of permutation matrices. Thus each swap of regions *k* and *l* in Π for which Ω_*kl*_ = 1 incurs a penalty cost of 2l. We set *λ* = 3 x 10^−4^ throughout, as this value was the smallest penalty obtained by a coarse grid search that suppressed all swaps among the penalized region-pairs in the 41 pairs of test-retest FCs (FCs from the same individual 6 months apart) for both versions of penalties.

To optimize (1), we used an iterative linear assignment procedure initialized at the identity Π^(0)^ = **I**, which represents the case where no swaps occur between the two matrices. This initialization amounts to implicit regularization toward the graph-matching solution with no swaps. Noting that 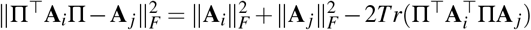 and 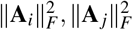 are fixed constants with respect to Π, (1) is equivalent to the following:

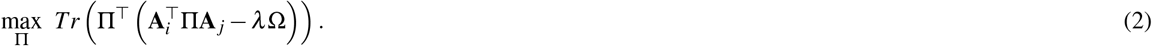

Exact optimization of (2) over the set of permutation matrices is generally intractable. We used instead the following iterative procedure:

1. Set 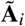 and 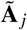 to be the matrices **A**_*i*_ and **A**_*j*_ with diagonal entries replaced by 0 and normalized to have Frobenius norm equal to 1.
2. Initializing at Π^(0)^ = **I**, for *t* = 1, 2,..., *t*_max_, repeat

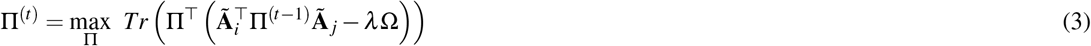

until the objective function (2) converges.

Setting the diagonals in the FC matrices to 0 ensures that regions’ self connections (which in the precision FCs used here are not all 1 as they are in the Pearson FC case) are not used to perform matching. Step 2 consists of iteratively solving a sequence of linear assignment problems. For this, we applied the classic Hungarian algorithm^50^ as implemented in the matchpairs() function in MATLAB. In general, this yields only a local maximum of the objective function (3). However, the attained objective value was consistently higher than an alternative state-of-the-art approach based on convex relaxation using quadratic programming, which seeks to obtain a more “global” optimum^51,52^. We believe this is because in our application, the optimal permutation in (3) should be close to the identity in most instances. Thus the initialization at Π^(0)^ = **I** renders this iterative approach more effective at finding a good local maximizer of (3), and also serves to regularize the final permutation matrix towards the identity.

In practice, we set *t*_max_ = 10 and run an iterative procedure until the difference in objective functions is smaller than 10^−3^. This corresponds to 4 or 5 iterations for the majority of pairs of individuals’ FCs.

### 2.3 Calculation of graph-matching based metrics

The optimal graph-matching permutation matrices Π_*i, j*_ as computed above have exactly one entry equal to 1 in each row or column, and 0 elsewhere.

An entry of 1 on the diagonal indicates that region’s FC in one individual *i* corresponds to the same region’s FC in the other individual *j*, whereas off-diagonal 1’s indicate that swapping those two different regions’ FC profiles results in the two subjects’ FCs being more similar. The permutation matrices can be analyzed in various ways – one can quantifying either the total number of off-diagonal swaps between subjects (as a measure of inter-subject differences) or the swap frequency of regions or networks across all pairs of subjects (as a measure of region’s/region-pair’s contribution to inter-subject differences). Formally,

- **Swap distance**, defined for pairs of individuals, is the percentage of off-diagonal entries in Π. This value represents the distance between FC matrices from subject *i* and subject *j*; lower values of swap distance indicate that pair of subjects’ FCs are more similar.
- **Swap frequency**, defined for regions or pairs of regions, quantifies how often a region or a pair of regions was swapped across all pairs of individuals. Regional and region-pair swap frequencies can also be summarized at a network or network-pair level.

All statistical analyses were performed using MATLAB. Results corresponding to the more strict graph-matching penalty, i.e. penalizing swaps between regions in the limbic, sub-cortical and cerebellar networks and any other region in the brain, are presented in the main text. Results corresponding to the less strict graph matching penalty, i.e. penalizing swaps between pairs of regions within limbic, subcortical and cerebellar networks only, are presented in the supplement.

### 2.4 Analysis of swap distance

We began by investigating swap distance across five categories of subject relatedness: self (test-retest data for 41 subjects), monozygotic twins (MZ, 130 pairs), dizygotic twins (DZ, 68 pairs), full siblings (FS, 200 pairs), unrelated individuals that were not age-matched (UR Non Age-Matched, 459923 pairs) and unrelated individuals that were age-matched (UR Age-matched, 36,177). To compare the swap distance between multiple categories of relatedness, we used one-way analysis of variance (ANOVA) followed by Student’s t-tests; p-values were corrected for multiple comparisons using Bonferroni correction^53^ and values less than or equal to 0.05 were considered significant. Unrelated pairs of individuals were separated into age-matched and not age-matched categories to account for any age effects in the relatedness analysis.

Next, we quantified associations between swap distance and the ages of the pairs of individuals that had ages within the range of 22-35; 9 individuals aged 36 and over were excluded due to small sample size. To assess the association between swap distance and age, we fit a simple linear model

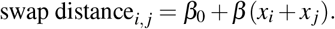

where *x_i_* and *x_j_* were the ages of subject *i* and *j*, respectively. We tested the significance of *β* > 0 using a permutation test, where ages were randomly permuted among individuals for 10,000 times to generate the empirical null distribution. This analysis was also conducted for female-female pairs and male-male pairs separately. Finally, we used a t-test to check for differences in swap distance between female-female and male-male pairs. For this, we used a subset of 200 unrelated, age-matched individuals (100 females and 100 males).

Finally, we tested whether the swap distance between pairs of individuals was related to their Total Cognition Scores, a measure of executive function derived from the NIH Toolbox Cognition battery^54–61^. Total cognition scores average the Crystallized Composite score, which seeks to measure skills acquired through life, and the Fluid Composite Score, which seeks to measure executive function/cognitive flexibility and episodic memory^62^. For this analysis, we had to exclude 11 subjects for whom cognitive scores were not available. We fit a linear model

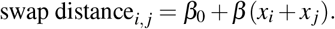

where *x_i_* and *x_j_* were the total cognition scores (CTC) of subject *i* and *j* to test the association between pairwise CTC and swap distance. CTC were also permuted 10,000 times to construct a empirical null distribution, upon which p-values were evaluated. This analysis was also conducted using swap distances of female-female pairs and male-male pairs separately.

### 2.5 Analysis of swap frequency

We used four ways of assessing the spatial distribution of swaps for a single individual: 1) region-pair swap frequencies, defined as the fraction of the 996 (other) individuals in which a swap occurred between that pair of regions, 2) regional swap frequencies, defined as the fraction of the 996 individuals that had a swap for that region (regardless of the other region involved in the swap) 3) network-pair swap frequencies, which are the region-pair swap frequencies averaged at the network-pair level and 4) network swap frequencies, which are the regional swap frequencies averaged at the network level. These four metrics were calculated for each individual and then summarized at a population-level by averaging them across all 997 subjects. Average network swap frequencies were compared using one-way analysis of variance (ANOVA) and student’s t-tests.

Finally, we calculated regional swap frequencies for pairs of individuals within the familial relatedness categories. To assess how the spatial distributions of swaps varies with relatedness, we calculated the Spearman rank correlation of each regions’ swap frequencies with relatedness coded as integers - 1 for self, 2 for monozygotic twins, 3 for dizygotic twins, 4 for full siblings and 5 for unrelated (not age-matched) individuals.

### 2.6 Code availability

The denoising, regional time series extraction, volume censoring, and FC calculation code is publicly available at https://github.kjamison/fmriclean. Codes for generating precision FC matrices can be found at https://github.com/zijin-gu/scfc-coupling. The code for graph matching is available through https://github.com/ChangSuBiostats/pairwise_graph_matching. Codes for swap percentage and frequency analyses can be found at https://github.com/hbukhari174/GM_HCP.

## 3 Results

### 3.1 Swap distance between pairs of individuals increases monotonically with their degree of relatedness

Swap distance between pairs of individuals varies with their degree of relatedness (Fig. 2A); one-way analysis of variance (ANOVA) revealed significant differences in swap distance by individuals’ relatedness (*p* = 0 after correction). The smallest swap distance occurred for within-individual test-retest pairs (self) wherein only 1.08% ± 0.78% of regions swapped with a different region, followed by 7.69% ± 0.44% for pairs of monozygotic (MZ) twins, 12.74% ± 0.60%) for pairs of dizygotic (DZ) twins, 12.98% ± 0.36%) for full siblings, 18.44% ± 0.026%) for age-matched unrelated individuals, and 18.40% ± 0.007%) for non-age matched unrelated individuals. Pairs of monozygotic (MZ) twins had a significantly higher average swap distance compared to self (*p* = 3.14*e* – 12 after correction), and significantly lower swap distance compared to 1) DZ twins (corrected *p* = 2.97*e* – 10), 2) full siblings (FS) (corrected *p* = 1.43*e* – 19) and 3) unrelated age-matched pairs (corrected *p* = 0) and 4) unrelated not age-matched pairs (corrected *p* = 0). Swap distance between pairs of DZ twins was not significantly different from that of full siblings, but both were significantly larger than test-retest self pairs (corrected *p* = 1.38*e* – 30, *p* = 3.35*e* – 42). Pairs of unrelated individuals (not age-matched and age-matched) did not have significantly different swap distance compared to each other, but both groups had significantly higher swap distance than test-retest self pairs (corrected *p* = 0), DZ twins (corrected *p* = 0) and FS pairs (corrected *p* = 0). The effect of the two penalty terms can be appreciated in Supplemental Figure S1 which shows the self test-retest region-pair swap frequency matrix for the no penalty case (A), less strict penalty case (B) and more strict penalty case (C).

**Figure 1.**
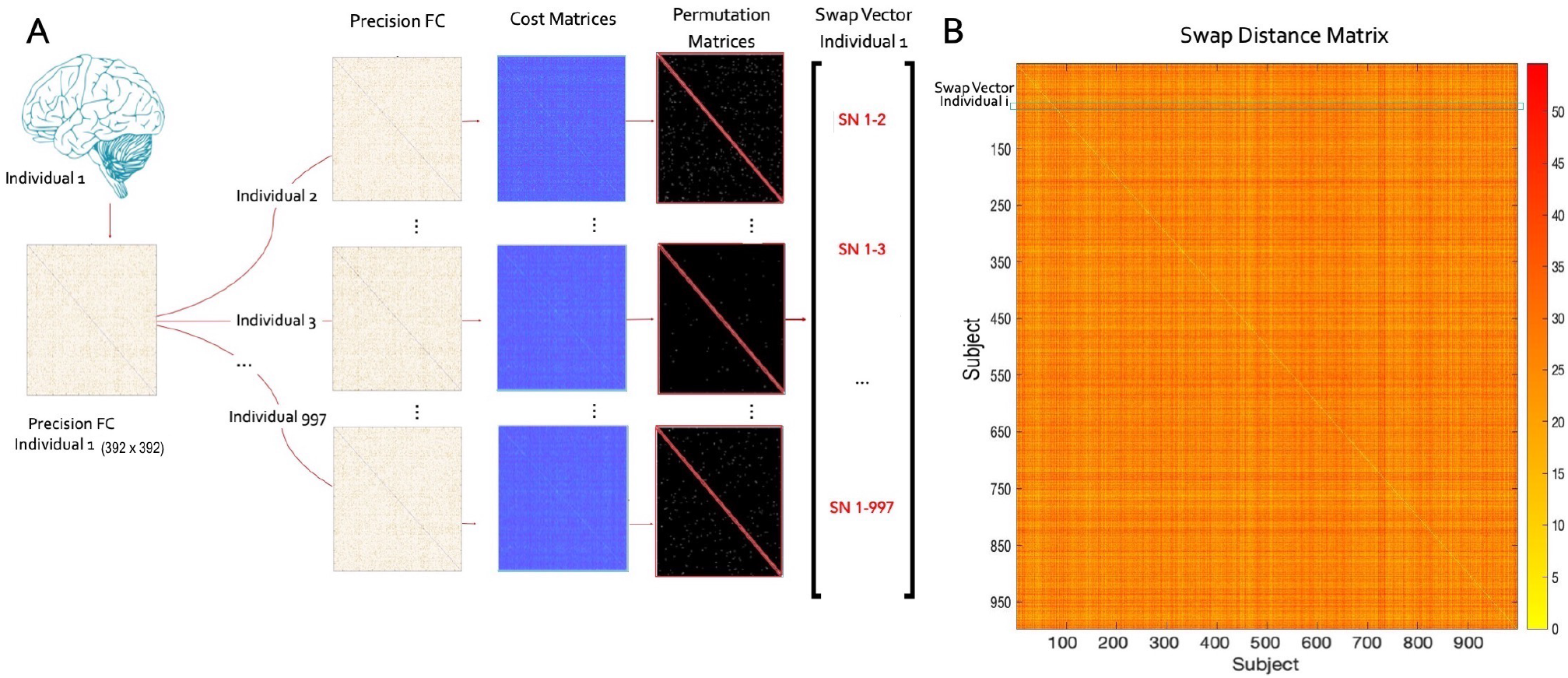
(**A**) Graph matching of functional connectomes (FCs). FCs are mapped between pairs of individuals such that rows (a single region’s connections to the rest of the brain) are permuted (swapped) with other rows in that individual’s FC if doing so minimizes the cost matrix (i.e. maximizes similarity to the other individual’s FC). The permutation matrix records swaps as 1’s on the off diagonal so that the sum of the off-diagonal elements represents the number of swapped regions; there are 996 permutation matrices and 996 sums of off-diagonal swaps per individual. (**B**) Population level results are represented by the 997 x 997 distance matrix containing swap distances between each pair of individuals; 0’s along the diagonal indicate no swapping between the same subject’s FC.

**Figure 2.**
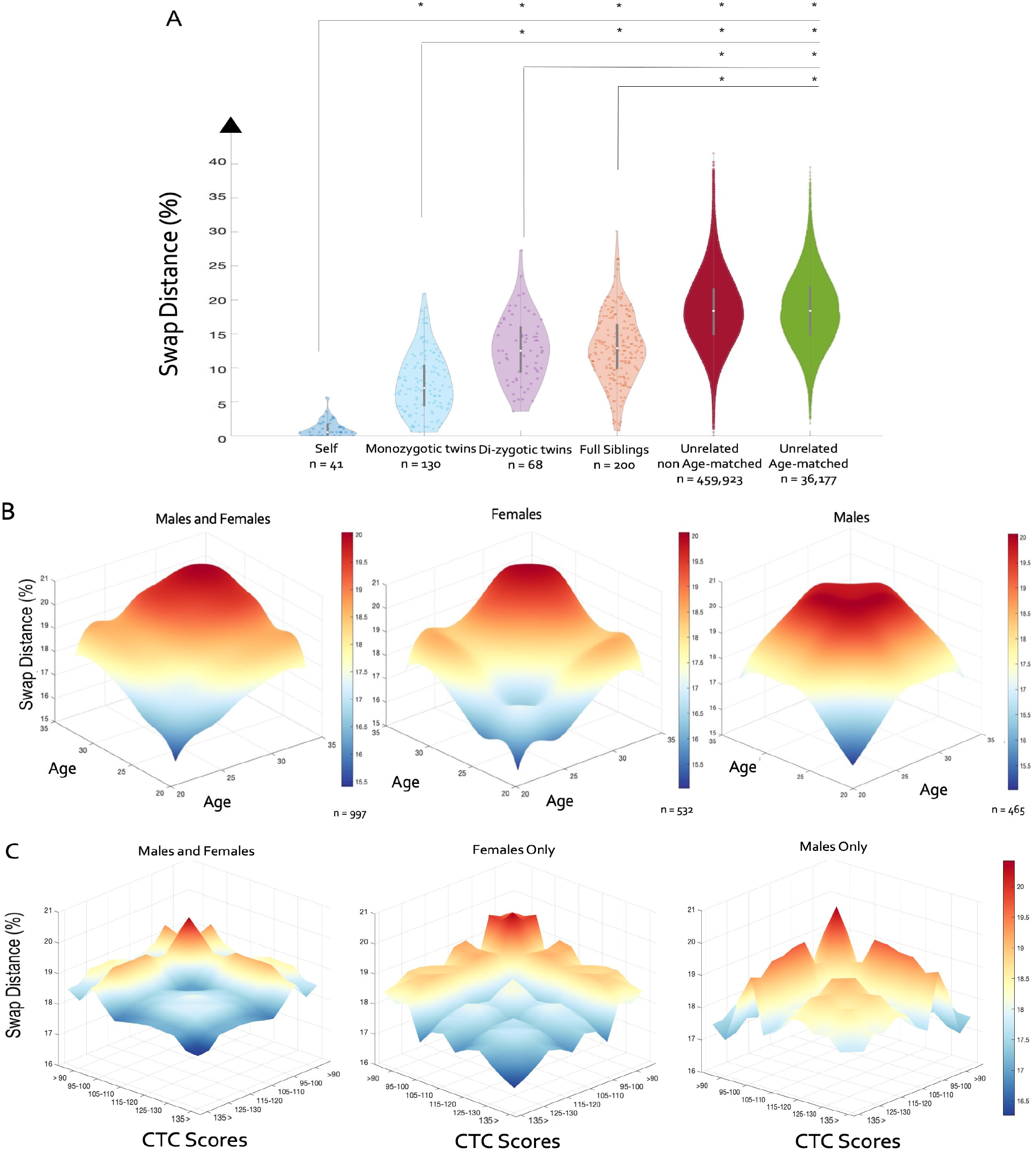
(A) Average swap distance between pairs of test-retest (self), mono-zygotic, di-zygotic, full siblings, un-related non age-matched and un-related age-matched subjects, * indicates significant differences in swap percent between the groups in question (corrected p<0.05). (B) Swap distance as a function of the age of the pairs of individuals; all pairs are illustrated on the left most panel, while only female-female pairs and male-male pairs are shown in the middle and right, respectively. (C) Swap distance between pairs of individuals indexed by total cognition (CTC) scores; all pairs of individuals regardless of sex are on the left, with female-female and male-male pairs are shown in the middle and right, respectively. Axes for CTC scores are reversed.

### 3.2 Swap distance increases with age and sex

We calculated the average swap distance for all possible unique pairs of individuals’ ages in our analysis (Figure 2B). Swap distance was largest when both individuals were from the higher end of the age range, decreased as at least one individual’s age decreased, and had the lowest value when both individuals were on the youngest end of the age range. Significant, positive associations between age and swap distance were identified via linear regression for all pairs of individuals (regardless of sex) (*p* = 0), pairs of females (*p* = 0) and pairs of males (*p* = 0). In our analysis of the 200 unrelated, age-matched individuals (100 male and 100 female), we found that the swap distance between pairs of females (17.46% ± 2.83) was smaller compared to the swap distance between pairs of males (19.38% ± 2.83) (*t* (198) = –4.91, *p* = 1.89*e* – 06).

### 3.3 Swap distance relates to individuals’ cognitive scores

The number of pairs of individuals with each pair of discretized cognitive scores is reported in Supplemental Figure S6. Swap distance appeared to vary with cognitive scores when considering both male and female individuals (Figure 2C)). However, when separated by sex, this apparent relationship persisted only in pairs of females wherein females with lower CTC scores had overall higher swap distances from every other female. Indeed, significant, negative associations between CTC and swap distance were identified via linear regression only for pairs of females (*p* = 0.0098) but not pairs of males (*p* = 0.75).

### 3.4 Regional and network-level swap frequencies

We found that association networks, namely fronto-parietal, default mode and ventral/dorsal attention, exhibited highest network-level swap frequencies, while sensory-motor networks exhibited lower overall swap frequencies, see Figure 3A and B. The highest within network-pair swap frequency occurred between pairs of regions that were both within FPN, DMN and ventral attention (VA) networks, while the highest across network swaps occurred between regions in the FPN and DMN (Note: the graph matching penalty suppressed most swaps to or from the limbic, sub-cortical and cerebellar networks.)

**Figure 3.**
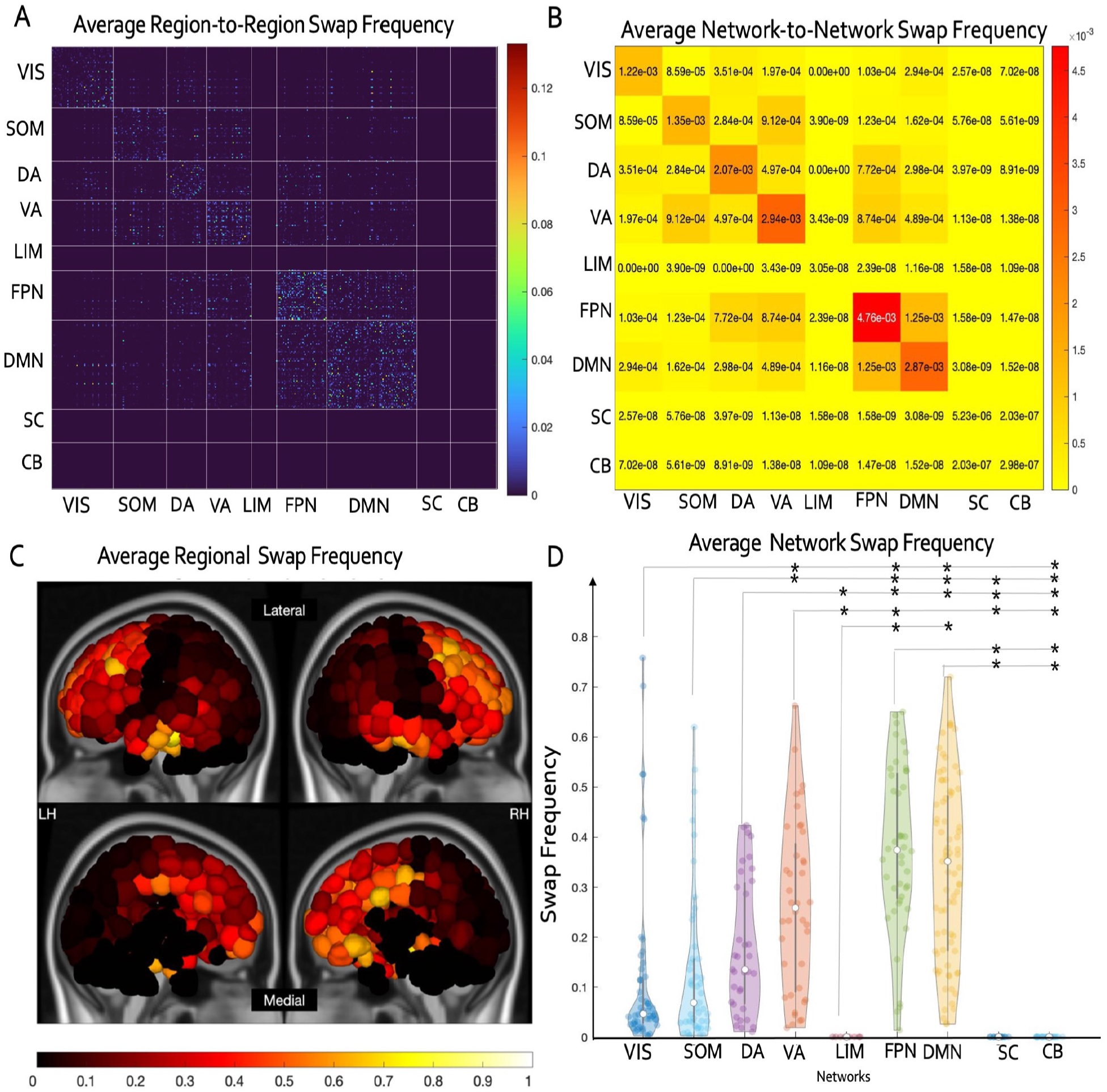
(A) Region-pair swap frequency and (B) network-pair swap frequencies, illustrating the frequency of swapping between pairs of regions and pairs of networks, respectively, averaged over all pairs of individuals. (C) Regional and (D) network-level summaries of swap frequencies, * indicates significant differences in network swap frequency (corrected p < 0.05). (Vis = visual, Som = somatomotor, DA = dorsal attention, VA = ventral attention, Lim = Limbic, FPN = Fronto-parietal, DMN = default mode, Cb = cerebellum, Sc= subcortical).

Networks showed significant differences in between their average network swap frequencies (One-way ANOVA, *p* = 9.79*e* – 40 after corrections for multiple comparisons) (Figure 3C and D). Regions from the fronto-parietal and default mode networks had the highest average network swap frequency (Figure 3C and D). There were no significant differences in network swap frequency between the visual and somatomotor networks, which exhibited lower swap frequencies in comparison to most other networks.

### 3.5 Regional swap frequencies correlate with degree of relatedness

Spearman rank correlation of relatedness (self, MZ, DZ, FS and unrelated, not age-matched) and regional swap frequency between pairs of individuals within the same relatedness category revealed that swap frequency increased monotonically with degree of relatedness for most regions in association networks, but not for as many regions in the somatomotor or visual networks (Fig. 4.)

**Figure 4.**
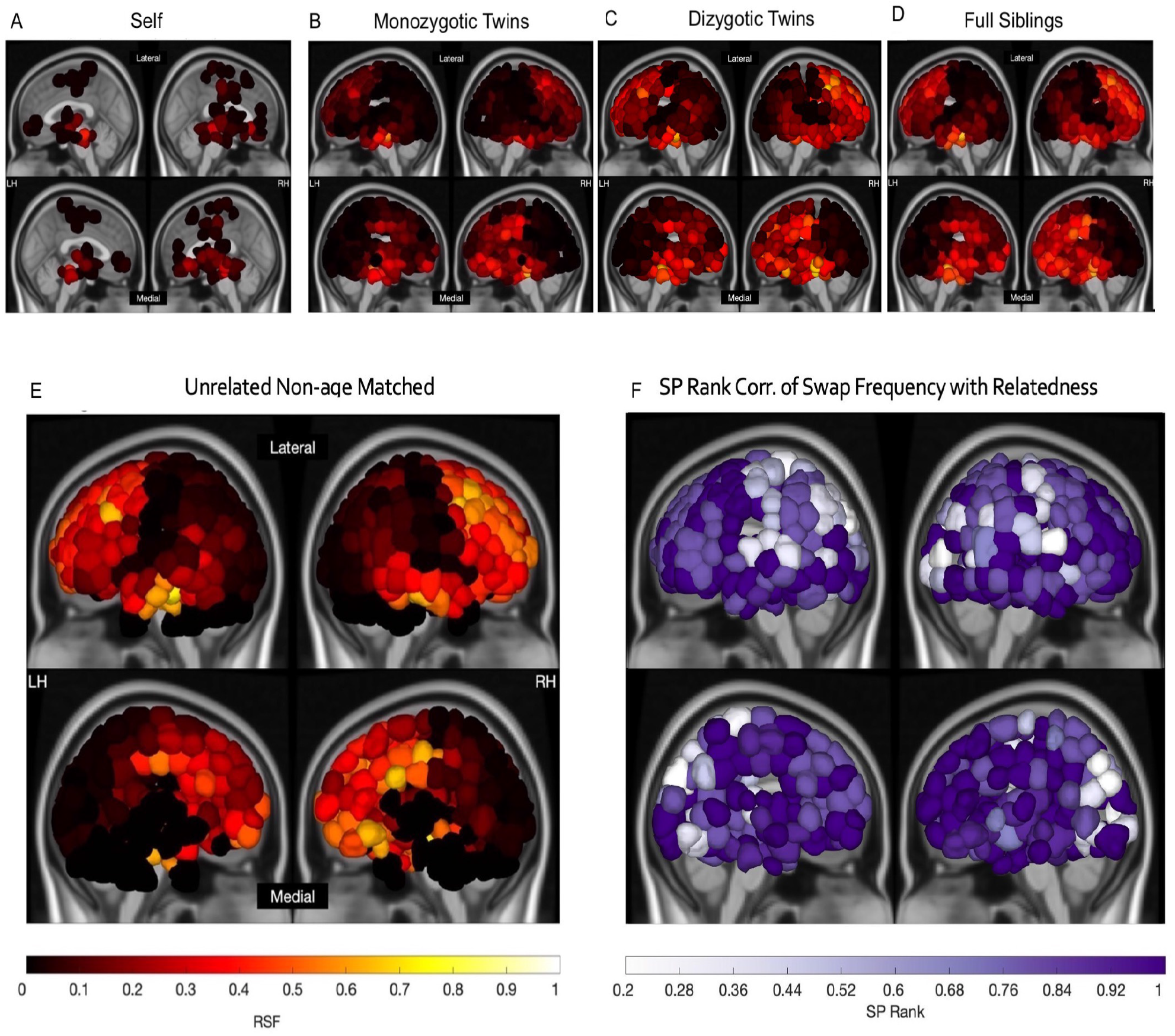
Average regional swap frequencies across pairs of individuals within the same relatedness category: A) self (test-retest), B) monozygotic twins (MZ), C) dizygotic twins (DZ), D) full siblings (FS) and E) unrelated, not age-matched (UR-Na). F) Each region’s Spearman’s rank correlation of the values in A-F (average regional swap frequency for each familial relatedness category) and an integer denoting level of familial relatedness. Here, values of 1 indicate a monotonically increasing average swap frequency with increasing familial relatedness distance.

All of these results are using graph matching with the more strict penalty for swaps involving at least one region from limbic, subcortical and cerebellar networks. However, the results were identical when using the less strict penalty that penalized swaps between pairs of regions that were both in the limbic, subcortical and cerebellar networks, see Supplemental Figures S4, S5 and S6.

## 4 Discussion

In this study, we used a novel graph-matching approach to quantify the similarity between pairs of individuals’ FCs in the large-scale HCP-YA dataset. We showed that the graph-matching metric between pairs of individuals, called swap distance, 1) increased monotonically with degree of relatedness (self < monozygotic < dizygotic = full siblings < unrelated individuals), 2) was largest when both individuals were on the higher end of the age range, decreased as one individual decreased in age and was smallest when both individuals were on the lower end of the age range, 3) was smaller in female pairs compared to male pairs and 4) may be related to total cognition score in females, wherein those females lower scores had overall more swaps than those with higher scores. We also showed that regional swap frequency varied with level of functional hierarchy: lower-order sensory-motor regions had the most similarity among pairs of individuals (lowest swap frequency) and higher-order association regions, particularly default-mode and fronto-parietal regions, had the least similarity among pairs of individuals (highest swap frequency). Finally, we showed that the swap frequency of association network regions, and not sensory-motor regions, varied monotonically with degree of relatedness between the pairs of individuals. This work demonstrates the utility of a novel graph matching metric for quantifying variability of whole-brain functional connectomes between pairs of individuals by showing it is associated with degree of relatedness, ages and cognitive scores between the pairs of individuals in question. We further demonstrate that the most variability across individuals is in higher-order association networks, i.e. the default-mode and fronto-parietal networks, that are associated with memory, emotional regulation and executive functioning.

The large majority of brain-behavior mapping literature rests on group-wise statistical tests or population-level regression models to map brain biomarkers to some demographic or cognitive outcome. Our study steps away from analysis performed at a group level or individual vs group level, and instead attempts to measure similarity at the level of pairs of individuals. There are a few other works that have investigated connectome similarities between pairs of individuals by computing correlations or cosine distances of individuals’ FCs in order to classify and identify connectomes across time or to see if connectomes are heritable^11,26,63^. Other individual-centring works have focused on temporally aligning time series with the goal of better identifying connectomes, generating individual parcel schemes, or categorizing disease-specific neural dynamics^34,35,64^. Investigating the pairwise differences among individuals as we do here allows a rich assessment of how age, sex, familial relatedness and cognitive scores may impact inter-subject variability in FCs.

While this paper is the first to apply graph matching to investigate the similarities of FCs between individuals, previous work has used graph matching to measure similarity between FC and SC and individuals’ SCs. One such work in individuals with traumatic brain injury (TBI) used graph matching to capture SC network integrity after injury and in recovery; specifically they showed that individuals with TBI have more heterogeneity in their SCs than healthy controls, and this heterogeneity increases over time and is related to cognitive scores^65^. Graph matching has also been used in the past to map SC and FC to one another; that study found highest structure-function alignment in visual and somatomotor networks and lowest alignment in association regions^43^, which reflects our current results showing least inter-subject FC alignment for association and most for sensory-motor networks. They also showed SC-FC alignment was similar for males and females and decreased with age across development. We conjecture the latter finding is linked our current results showing increased swap distance with increasing age - if there is less alignment of SC and FC, then there may be more divergence of FC across the population. In our recent work, the same graph-matching approach was used to track longitudinal FC reorganization in individuals with first-time pontine stroke^41^. There, we used graph matching to assess swap distance between the same individual’s FCs over the post-stroke recovery period. We showed that brain regions with more structural (white matter) disconnection and functional connectivity disruption due to the stroke lesion had more swapping of FC profiles over time, and, importantly, that individuals with more regional swaps over time also had better recovery of motor function. There, we proposed our graph-matching approach as a way to track recovery-relevant functional reorganization over time in the same individual, while here we use it to quantify inter-individual differences in FC patterns.

In alignment with many other works confirming the heritability of FC, we identified a monotonically increasing swap distance as degree of familial relatedness increased^11,16,18,66–68^. The lowest swap percent for test-retest (self) pairs reflects the relative stability inherent to a person’s connectome over time, while the difference between test-retest pairs and monozygotic twins indicates the potential role of the environment or personal experience in shaping the functional organization of the brain, particularly for higher-order default mode and fronto-parietal regions^10,11,28,69^. Moreover, the significant difference between monozygotic twins and dizygotic twins/full siblings (who presumably all share a similar environment) suggests the impact that genetics can play on FC patterns.

FC has been shown to change with development and aging, even across the relatively restricted period of early adulthood^70–74^. Throughout the young adult age range studied here, the brain faces a variety of structural changes, including an increase in ventricle size and loss of gray and white matter volume in different cortical areas^75^. Across development, alterations to the brain lead different networks to show both gain and deficit in distinctive FC patterns that manifest at a local and global level^70^. Both higher and lower order networks like the attention, limbic or visual show fluctuations in FC within and between networks, pointing to the diversity of FC patterns and/or their global topologies as we continue to age^76^. In our work, we find that swap distance between pairs of individuals increases as one or both of their ages increase, regardless of their sex. This indicates that there is generally a divergence of the FC patterns as individuals in early adulthood age. The relationship between FC and cognition can be influenced by a variety of environmental factors ranging from education level to socioeconomic status, sensitive to changes in age, can be domain/task-specific and exhibits sex-specific variability?’^48,77–81^. Here we identified a larger swap distance for female subjects with lower total cognition scores compared to those with higher scores; we conjecture this could be reflective of a number of different environmental, experiential or biological factors which extend beyond the scope of this study. Finally, using a subset of 100 age-matched males and 100 age-matched females (all unrelated), we found that swap distance was smaller among female-female pairs compared to male-male pairs. This aligns with previous work identifying less inter-subject variance in females compared to males; however this effect could also be explained by other physiological factors, like head size, that we did not control for in this study^37,82–85^

We find that higher order networks, i.e. default mode and fronto-parietal, which underlie complex cognition, self-referential memory and executive function, are the most different across individuals aligns with previous work identifying that FC within higher order networks drives the classification and identification of individual connectomes^11,26,28,69^. We further found that regional swap frequency between individuals within the familial relatedness groups unsurprisingly had a monotonically increasing relationship with the degree of relatedness of that group, but only for regions in the association networks that had highest overall swap frequencies and were presumably driving the individual-pair swap distances. This result further validates the regional swap frequency metric is not just measuring noise, and that this region-specific metric (at least in association networks) may also be related to genetics/environment.

This analysis has some limitations. While we use a relatively large data set, the age range of the HCP-YA is rather limited and we do not validate our findings with out of site data. In the future, we aim to validate our findings with data from other sites and across different image processing pipelines, development, aging, behavioral/cognitive measures and genetic relatedness. Here we added a penalty to suppress self-to-self swaps to and from regions with lower SNR, including limbic, subcortical and cerebellar regions, which limits interpretation of the results from these areas. Future work using higher SNR fMRI data could allow interpretation of swap frequencies in these networks. Our analysis for age and total cognition also exclude swaps from pairs above the age of 35 and people scoring a CTC score over to 145 avoid over-representation due to low number of subjects.

Using the HCP-YA dataset, we quantify the variability of FCs across a diverse set of individuals using a metric defined at the level of pairs of individuals. We show that this metric is related to the individual-pairs’ degree of relatedness, ages and, only in females, cognitive abilities. With the largest swap frequency occurring in higher-order networks like the default-mode and fronto-parietal network, i.e. networks associated with memory, reward signalling and executive functioning, our results possibly reflect a combination of each individual’s lived experience, genetics and physiology.

## Acknowledgements

We would like to sincerely thank Dr. Zhou Fan at Yale University for his many helpful discussions on graph matching and its application to this data. This work was funded by the following grants: R01 NS102646 (A.K.), RF1 MH123232 (A.K.), and R21 NS120227-01 (A.K.).

## Notes

### Competing Interest Statement

The authors have declared no competing interest.

## References

1. Sporns, O., Tononi, G. & Kötter, R. The human connectome: a structural description of the human brain. PLoS computational biology 1, e42 (2005).

2. Varoquaux, G. & Craddock, R. C. Learning and comparing functional connectomes across subjects. NeuroImage 80, 405–415 (2013).

3. Van Essen, D. C. et al. The wu-minn human connectome project: an overview. Neuroimage 80, 62–79 (2013).

4. Smith, S. M. et al. Functional connectomics from resting-state fmri. Trends cognitive sciences 17, 666–682 (2013).

5. Smith, S. M. et al. Resting-state fmri in the human connectome project. Neuroimage 80, 144–168 (2013).

6. Cole, M. W., Bassett, D. S., Power, J. D., Braver, T. S. & Petersen, S. E. Intrinsic and task-evoked network architectures of the human brain. Neuron 83, 238–251 (2014).

7. Miller, K. L. et al. Multimodal population brain imaging in the uk biobank prospective epidemiological study. Nature neuroscience 19, 1523–1536 (2016).

8. Haimovici, A., Tagliazucchi, E., Balenzuela, P. & Laufs, H. On wakefulness fluctuations as a source of bold functional connectivity dynamics. Scientific reports 7, 1–13 (2017).

9. Power, J. D., Plitt, M., Laumann, T. O. & Martin, A. Sources and implications of whole-brain fmri signals in humans. Neuroimage 146, 609–625 (2017).

10. Ge, T., Holmes, A. J., Buckner, R. L., Smoller, J. W. & Sabuncu, M. R. Heritability analysis with repeat measurements and its application to resting-state functional connectivity. Proceedings National Academy Sciences 114, 5521–5526, DOI: 10.1073/pnas.1700765114 (2017). Doi: 10.1073/pnas.1700765114.

11. Miranda-Dominguez, O. et al. Heritability of the human connectome: A connectotyping study. Network Neuroscience 2, 175–199 (2018).

12. Dubois, J., Galdi, P., Paul, L. K. & Adolphs, R. A distributed brain network predicts general intelligence from resting-state human neuroimaging data. Philosophical Transactions Royal Society B: Biological Sciences 373, 20170284 (2018).

13. Power, J. D. et al. Distinctions among real and apparent respiratory motions in human fmri data. NeuroImage 201, 116041 (2019).

14. Elliott, M. L. et al. General functional connectivity: Shared features of resting-state and task fmri drive reliable and heritable individual differences in functional brain networks. Neuroimage 189, 516–532 (2019).

15. Lynch, C. J. et al. Prevalent and sex-biased breathing patterns modify functional connectivity mri in young adults. Nature communications 11, 1–14 (2020).

16. Barber, A. D., Hegarty, C. E., Lindquist, M. & Karlsgodt, K. H. Heritability of functional connectivity in resting state: Assessment of the dynamic mean, dynamic variance, and static connectivity across networks. Cerebral Cortex 31, 2834–2844 (2021).

17. Elam, J. S. et al. The human connectome project: A retrospective. NeuroImage 244, 118543 (2021).

18. Gu, Z., Jamison, K. W., Sabuncu, M. R. & Kuceyeski, A. Heritability and interindividual variability of regional structurefunction coupling. Nature Communications 12, 1–12 (2021).

19. Bullmore, E. & Sporns, O. Complex brain networks: graph theoretical analysis of structural and functional systems. Nature reviews neuroscience 10, 186–198 (2009).

20. Bassett, D. S. & Bullmore, E. T. Human brain networks in health and disease. Current opinion neurology 22, 340 (2009).

21. Bullmore, E. T. & Bassett, D. S. Brain graphs: graphical models of the human brain connectome. Annual review clinical psychology 7, 113–140 (2011).

22. Tompson, S. H., Falk, E. B., Vettel, J. M. & Bassett, D. S. Network approaches to understand individual differences in brain connectivity: opportunities for personality neuroscience. Personality neuroscience 1 (2018).

23. Craddock, R. C., James, G. A., Holtzheimer III, P. E., Hu, X. P. & Mayberg, H. S. A whole brain fmri atlas generated via spatially constrained spectral clustering. Human brain mapping 33, 1914–1928 (2012).

24. Wig, G. S. et al. Parcellating an individual subject’s cortical and subcortical brain structures using snowball sampling of resting-state correlations. Cerebral cortex 24, 2036–2054 (2014).

25. Gordon, E. M. et al. Precision functional mapping of individual human brains. Neuron 95, 791–807 (2017).

26. Finn, E. S. et al. Functional connectome fingerprinting: identifying individuals using patterns of brain connectivity. Nature neuroscience 18, 1664–1671 (2015).

27. Finn, E. S.et al. Can brain state be manipulated to emphasize individual differences in functional connectivity? Neuroimage 160, 140–151 (2017).

28. Horien, C., Shen, X., Scheinost, D. & Constable, R. T. The individual functional connectome is unique and stable over months to years. Neuroimage 189, 676–687 (2019).

29. Noble, S., Scheinost, D. & Constable, R. T. A decade of test-retest reliability of functional connectivity: A systematic review and meta-analysis. Neuroimage 203, 116157 (2019).

30. Bari, S., Amico, E., Vike, N., Talavage, T. M. & Goñi, J. Uncovering multi-site identifiability based on resting-state functional connectomes. NeuroImage 202, 115967 (2019).

31. Miranda-Dominguez, O. et al. Connectotyping: model based fingerprinting of the functional connectome. PloS one 9, e111048 (2014).

32. Greene, D. J. et al. Integrative and network-specific connectivity of the basal ganglia and thalamus defined in individuals. Neuron 105, 742–758 (2020).

33. Yao, D. et al. A mutual multi-scale triplet graph convolutional network for classification of brain disorders using functional or structural connectivity. IEEE transactions on medical imaging 40, 1279–1289 (2021).

34. Joshi, A. A., Chong, M., Li, J., Choi, S. & Leahy, R. M. Are you thinking what i’m thinking? synchronization of resting fmri time-series across subjects. NeuroImage 172, 740–752 (2018).

35. Joshi, A. A., Choi, S., Li, J., Akrami, H. & Leahy, R. M. A pairwise approach for fmri group studies using the brainsync transform. In Medical Imaging 2021: Image Processing, vol. 11596, 96–102 (SPIE, 2021).

36. Li, Y., Saxe, R. & Anzellotti, S. Intersubject mvpd: Empirical comparison of fmri denoising methods for connectivity analysis. PloS one 14, e0222914 (2019).

37. Hassanzadeh, R. et al. Individualized spatial network predictions using siamese convolutional neural networks: A resting-state fmri study of over 11,000 unaffected individuals. PloS one 17, e0249502 (2022).

38. Schellewald, C., Roth, S. & Schnörr, C. Evaluation of a convex relaxation to a quadratic assignment matching approach for relational object views. Image Vision Computing 25, 1301–1314 (2007).

39. Oyarzun Laura, C., Wesarg, S. & Sakas, G. Graph matching survey for medical imaging: On the way to deep learning. Methods (San Diego, Calif.) (2021).

40. Osmanlioğlu, Y., Alappatt, J. A., Parker, D. & Verma, R. Connectomic consistency: a systematic stability analysis of structural and functional connectivity. J Neural Eng 17, 045004, DOI: 10.1088/1741-2552/ab947b (2020). Osmanlioğlu, Yusuf Alappatt, Jacob A Parker, Drew Verma, Ragini 2020/5/20.

41. Olafson, E. R. et al. Functional connectome reorganization relates to post-stroke motor recovery and structural and functional disconnection. NeuroImage 245, 118642 (2021).

42. Shen, R. S. et al. Graph matching based connectomic biomarker with learning for brain disorders. In Uncertainty for Safe Utilization of Machine Learning in Medical Imaging, and Graphs in Biomedical Image Analysis, 131–141 (Springer, 2020).

43. Osmanlioğlu, Y. et al. System-level matching of structural and functional connectomes in the human brain. Neuroimage 199, 93–104, DOI: 10.1016/j.neuroimage.2019.05.064 (2019). Osmanlioğlu, Yusuf Tunç, Birkan Parker, Drew Elliott, Mark A Baum, Graham L Ciric, Rastko Satterthwaite, Theodore D Gur, Raquel E Gur, Ruben C Verma, Ragini 2019/5/30.

44. Glasser, M. F. et al. The minimal preprocessing pipelines for the human connectome project. Neuroimage 80, 105–124 (2013).

45. Griffanti, L. et al. Ica-based artefact removal and accelerated fmri acquisition for improved resting state network imaging. Neuroimage 95, 232–247 (2014).

46. Salimi-Khorshidi, G. et al. Automatic denoising of functional mri data: combining independent component analysis and hierarchical fusion of classifiers. Neuroimage 90, 449–468 (2014).

47. Thomas Yeo, B. et al. The organization of the human cerebral cortex estimated by intrinsic functional connectivity. Journal neurophysiology 106, 1125–1165 (2011).

48. Dhamala, E., Jamison, K. W., Jaywant, A., Dennis, S. & Kuceyeski, A. Distinct functional and structural connections predict crystallised and fluid cognition in healthy adults. Hum Brain Mapp DOI: 10.1002/hbm.25420 (2021). 1097–0193 Dhamala, Elvisha Orcid: 0000-0002-8253-6962 Jamison, Keith W Orcid: 0000-0001-7139-6661 Jaywant, Abhishek Orcid: 0000-0003-4307-5742 Dennis, Sarah Orcid: 0000-0003-0370-4284 Kuceyeski, Amy Orcid: 0000-0002-5050-8342 R01NS102646-01A1/NH/NIH HHS/United States R21NS104634-01/NH/NIH HHS/United States 1K12-HD093427-04/National Center of Medical Rehabilitation Research/ Eunice Kennedy Shriver National Institute of Child Health and Human Development/ Journal Article United States Hum Brain Mapp. 2021 Apr 8. doi: 10.1002/hbm.25420.

49. Liégeois, R., Santos, A., Matta, V., Van De Ville, D. & Sayed, A. H. Revisiting correlation-based functional connectivity and its relationship with structural connectivity. Network Neuroscience 4, 1235–1251 (2020).

50. Kuhn, H. W. The Hungarian method for the assignment problem. Naval research logistics quarterly 2, 83–97 (1955).

51. Zaslavskiy, M., Bach, F. & Vert, J.-P. A path following algorithm for the graph matching problem. IEEE Transactions on Pattern Analysis Machine Intelligence 31, 2227–2242 (2008).

52. Aflalo, Y., Bronstein, A. & Kimmel, R. On convex relaxation of graph isomorphism. Proceedings National Academy Sciences 112, 2942–2947 (2015).

53. Weisstein, E. W. Bonferroni correction. https://mathworld.wolfram.com/ (2004).

54. Carlozzi, N. E. et al. Construct validity of the nih toolbox cognition battery in individuals with stroke. Rehabilitation psychology 62, 443 (2017).

55. Gershon, R. C. et al. Nih toolbox for assessment of neurological and behavioral function. Neurology 80, S2–S6 (2013).

56. Heaton, R. K. et al. Reliability and validity of composite scores from the nih toolbox cognition battery in adults. Journal International Neuropsychological Society 20, 588–598 (2014).

57. Mungas, D. et al. Factor structure, convergent validity, and discriminant validity of the nih toolbox cognitive health battery (nihtb-chb) in adults. Journal International Neuropsychological Society 20, 579–587 (2014).

58. Tulsky, D. S. et al. Factor structure of the nih toolbox cognition battery in individuals with acquired brain injury. Rehabilitation psychology 62, 435 (2017).

59. Weintraub, S. et al. I. nih toolbox cognition battery (cb): introduction and pediatric data. Monographs Society for Research Child Development 78, 1–15 (2013).

60. Weintraub, S. et al. The cognition battery of the nih toolbox for assessment of neurological and behavioral function: validation in an adult sample. Journal International Neuropsychological Society 20, 567–578 (2014).

61. Zelazo, P. D. et al. Nih toolbox cognition battery (cb): validation of executive function measures in adults. Journal International Neuropsychological Society 20, 620–629 (2014).

62. Heaton, R. K. et al. Reliability and validity of composite scores from the nih toolbox cognition battery in adults. Journal International Neuropsychological Society 20, 588–598 (2014).

63. Menon, S. S. & Krishnamurthy, K. A comparison of static and dynamic functional connectivities for identifying subjects and biological sex using intrinsic individual brain connectivity. Scientific reports 9, 1–11 (2019).

64. Kong, R. et al. Spatial topography of individual-specific cortical networks predicts human cognition, personality, and emotion. Cerebral cortex 29, 2533–2551 (2019).

65. Osmanlioğlu, Y. et al. Connectomic assessment of injury burden and longitudinal structural network alterations in moderate-to-severe traumatic brain injury. Human Brain Mapping 43, 3944–3957, DOI: https://doi.org/10.1002/hbm.25894(2022). https://onlinelibrary.wiley.com/doi/pdf/10.1002/hbm.25894.

66. Glahn, D. et al. Genetic control over the resting brain. Proceedings National Academy Sciences 107, 1223–1228 (2010).

67. Xu, J. et al. Heritability of the effective connectivity in the resting-state default mode network. Cerebral cortex 27, 5626–5634 (2017).

68. Demeter, D. V. et al. Functional connectivity fingerprints at rest are similar across youths and adults and vary with genetic similarity. Iscience 23, 100801 (2020).

69. Jalbrzikowski, M. et al. Functional connectome fingerprinting accuracy in youths and adults is similar when examined on the same day and 1.5-years apart. Human brain mapping 41, 4187–4199 (2020).

70. Tamnes, C. K. et al. Brain development and aging: overlapping and unique patterns of change. Neuroimage 68, 63–74 (2013).

71. Chan, M. Y., Park, D. C., Savalia, N. K., Petersen, S. E. & Wig, G. S. Decreased segregation of brain systems across the healthy adult lifespan. Proceedings National Academy Sciences 111, E4997–E5006 (2014).

72. Geerligs, L., Renken, R. J., Saliasi, E., Maurits, N. M. & Lorist, M. M. A brain-wide study of age-related changes in functional connectivity. Cerebral cortex 25, 1987–1999 (2015).

73. Zhang, C. et al. Sex and age effects of functional connectivity in early adulthood. Brain connectivity 6, 700–713 (2016).

74. Váša, F. et al. Conservative and disruptive modes of adolescent change in human brain functional connectivity. Proceedings National Academy Sciences 117, 3248–3253 (2020).

75. Sowell, E. R., Thompson, P. M. & Toga, A. W. Mapping changes in the human cortex throughout the span of life. The Neuroscientist 10, 372–392 (2004).

76. Betzel, R. F. et al. Changes in structural and functional connectivity among resting-state networks across the human lifespan. Neuroimage 102, 345–357 (2014).

77. Van Den Heuvel, M. P., Stam, C. J., Kahn, R. S. & Pol, H. E. H. Efficiency of functional brain networks and intellectual performance. Journal Neuroscience 29, 7619–7624 (2009).

78. Douw, L. et al. Cognition is related to resting-state small-world network topology: an magnetoencephalographic study. Neuroscience 175, 169–177 (2011).

79. Szalkai, B., Varga, B. & Grolmusz, V. Graph theoretical analysis reveals: Women’s brains are better connected than men’s. PLoS One 10, e0130045 (2015).

80. Pezoulas, V. C., Zervakis, M., Michelogiannis, S. & Klados, M. A. Resting-state functional connectivity and network analysis of cerebellum with respect to iq and gender. Frontiers human neuroscience 11, 189 (2017).

81. Dhamala, E., Jamison, K. W., Jaywant, A. & Kuceyeski, A. Shared functional connections within and between cortical networks predict cognitive abilities in adult males and females. Human brain mapping (2021).

82. Gur, R. C. et al. Sex differences in brain gray and white matter in healthy young adults: correlations with cognitive performance. Journal Neuroscience 19, 4065–4072 (1999).

83. Cosgrove, K. P., Mazure, C. M. & Staley, J. K. Evolving knowledge of sex differences in brain structure, function, and chemistry. Biological psychiatry 62, 847–855 (2007).

84. Forde, N. J. et al. Sex differences in variability of brain structure across the lifespan. Cerebral Cortex 30, 5420–5430 (2020).

85. Wierenga, L. M. et al. Greater male than female variability in regional brain structure across the lifespan. Human brain mapping 43, 470–499 (2022).

