## Supplemental Figures for "A graph-matching based metric of functional connectome distance between pairs of individuals varies with their ages, cognitive performances and familial relationships"

### ABSTRACT

Functional connectomes (FCs), represented by networks or graphs that summarize coactivation patterns between pairs of brain regions, have been related at a population level to age, sex, cognitive/behavioral scores, life experience, genetics and disease/disorders. However, quantifying FC differences between pairs of individuals also provides a rich source of information with which to map to differences in those individuals' biology, experience, genetics or behavior. In this study, graph matching is used to create a novel inter-individual FC metric, called swap distance, that quantifies the distance between pairs of individuals' FCs. We apply graph matching to align FCs between pairs of individuals from the the Human Connectome Project (N = 997) and find that swap distance i) increases with increasing familial distance, ii) increases with subjects' ages, iii) is smaller for pairs of females compared to pairs of males, and iv) is larger for females with lower cognitive scores compared to females with larger cognitive scores. Regions that contributed most to individuals' swap distances were in higher-order networks, i.e. default-mode and fronto-parietal, that underlie executive function and memory. These higher-order networks' regions also had swap frequencies that varied monotonically with familial relatedness of the individuals in question. We posit that the proposed graph matching technique provides a novel way to study inter-subject differences in FC and enables quantification of how FC may vary with age, relatedness, sex and behavior.

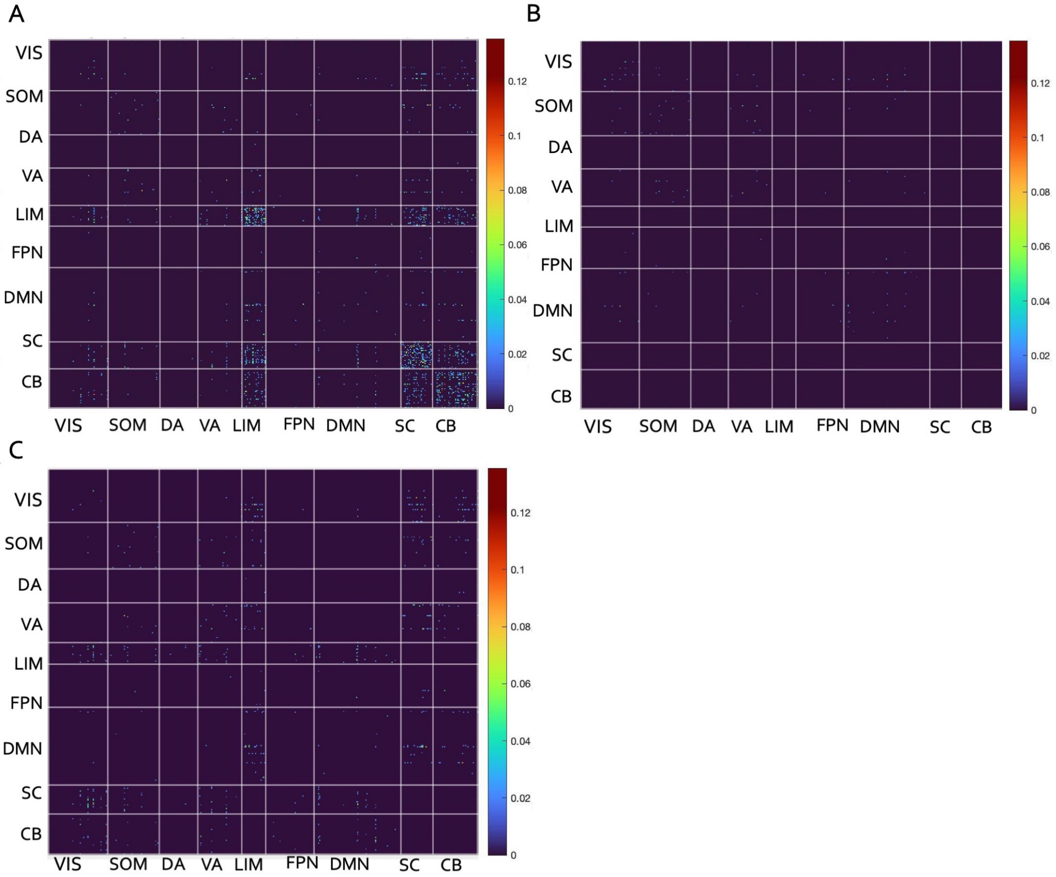


**Figure S1.** Region-pair swap frequencies calculated for test-retest pairs with no, the stricter and less strict penalty. (A) When no penalty applied in graph matching, region-pair swap frequencies for test-retest pairs were high within and between the limbic, subcortical, and cerebellar networks. (B) Region-pair swap frequencies for test-retest pairs when the penalty was applied to any swap involving limbic, subcortical and cerebellar regions and (C) and when the penalty was applied to swaps between pairs of regions in these three networks.

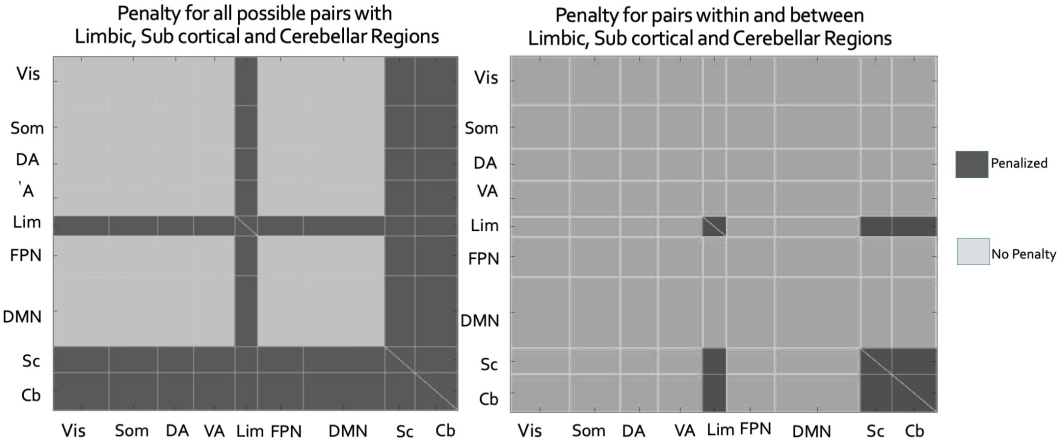


**Figure S2.** The stricter penalty was applied to any swap involving limbic, subcortical and cerebellar regions, while the less strict penalty was applied to swaps between pairs of regions in these three networks.

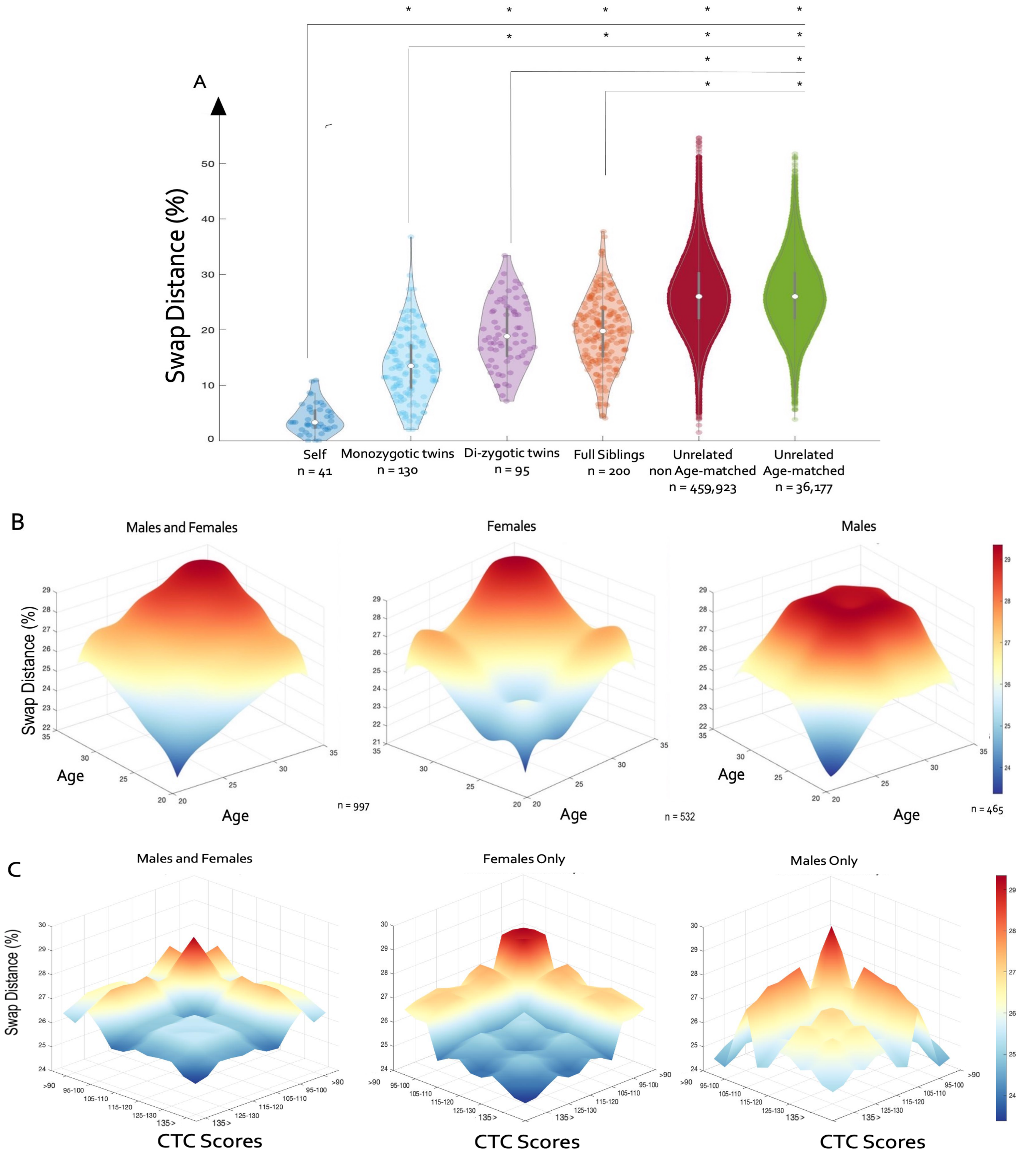


**Figure S3.** Swap distances calculated with the less strict penalty (penalizing swaps between pairs where both regions are in limbic, subcortical and cerebellar regions. (A) Average swap distance between pairs of test-retest (self), mono-zygotic (MZ), di-zygotic (DZ), full siblings (FS), un-related (UR) non age-matched and age-matched subjects. *indicates significant differences in swap percent between the groups in question (corrected p<0.05). (B) Swap distance as a function of the age of the pairs of individuals; all pairs are illustrated on the left most panel, while only female-female pairs and male-male pairs are shown in the middle and right, respectively. (C) Swap distance between pairs of individuals indexed by CTC scores; all pairs of individuals regardless of sex are on the right, with female-female and male-male pairs are shown in the middle and right, respectively.

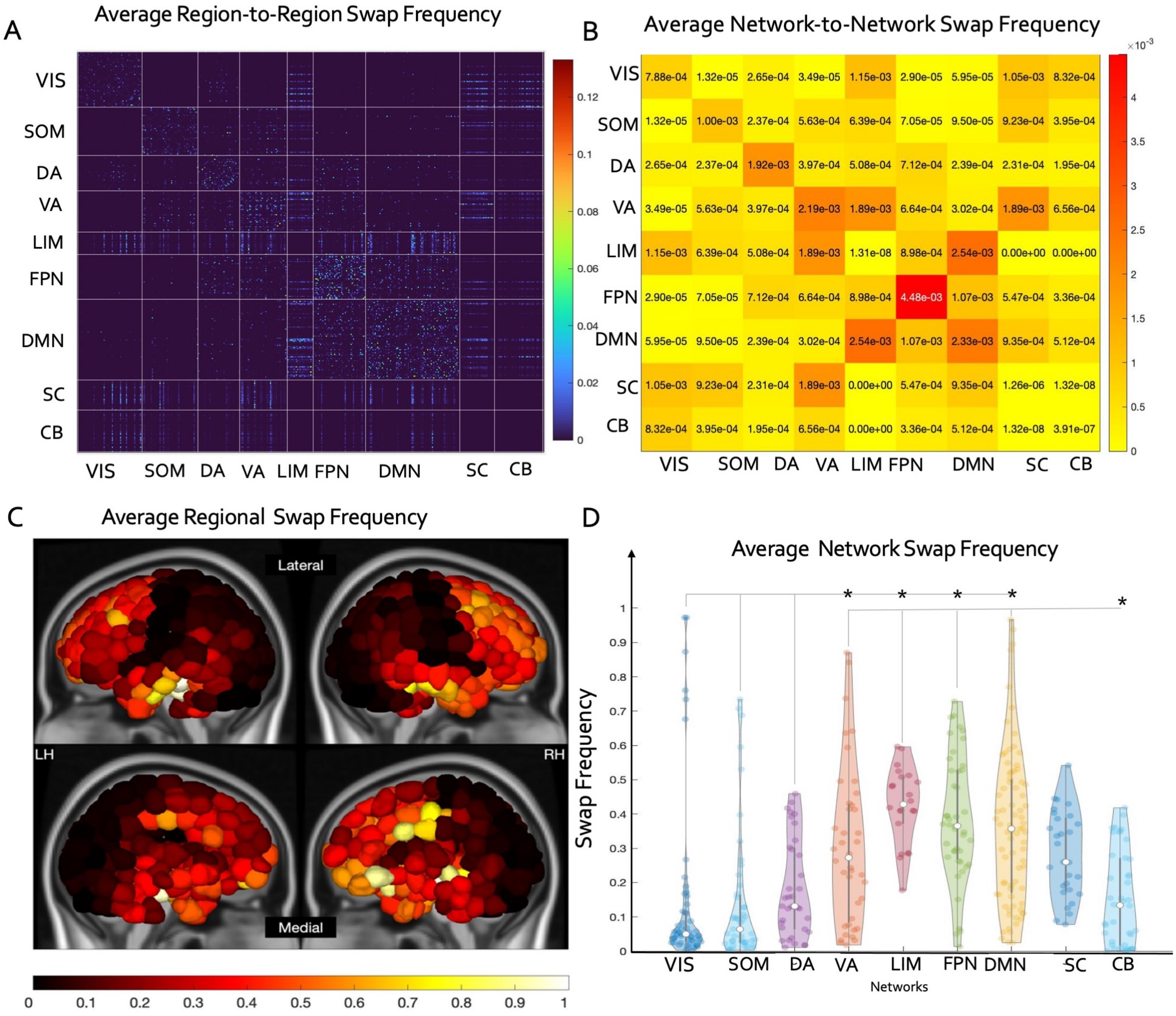


**Figure S4.** (A) Average region-pair swap frequency and (B) network-pair swap frequencies, (C) regional and (D) network-level summaries of swap frequencies with swapping penalized only between pairs of regions in limbic, subcortical and cerebellar networks. *indicates significant differences in network swap frequency (corrected p<0.05). (Vis = visual, Som = somatomotor, DA = dorsal attention, VA = ventral attention, Lim = Limbic, FPN = Fronto-parietal, DMN = default mode, Cb = cerebellum, Sc= subcortical).

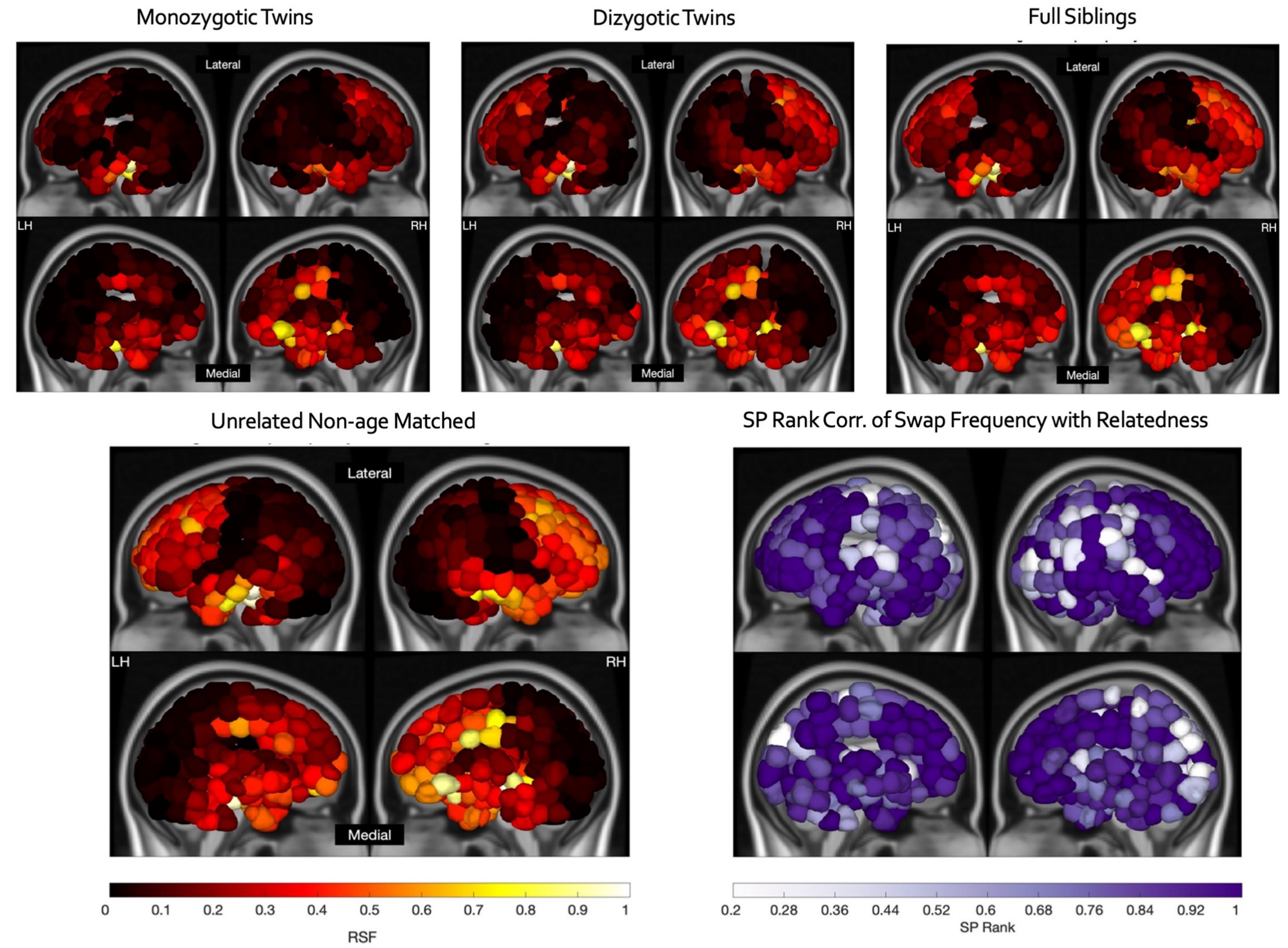


**Figure S5.** Average regional swap frequency was calculated for four categories of familial relatedness with swapping penalized only between pairs of regions in limbic, subcortical and cerebellar networks. (F) Spearman rank correlations between average regional swap frequency vs level of familial relatedness, i.e. self, followed by MZ, DZ, FS and, finally, UR (non age-matched).

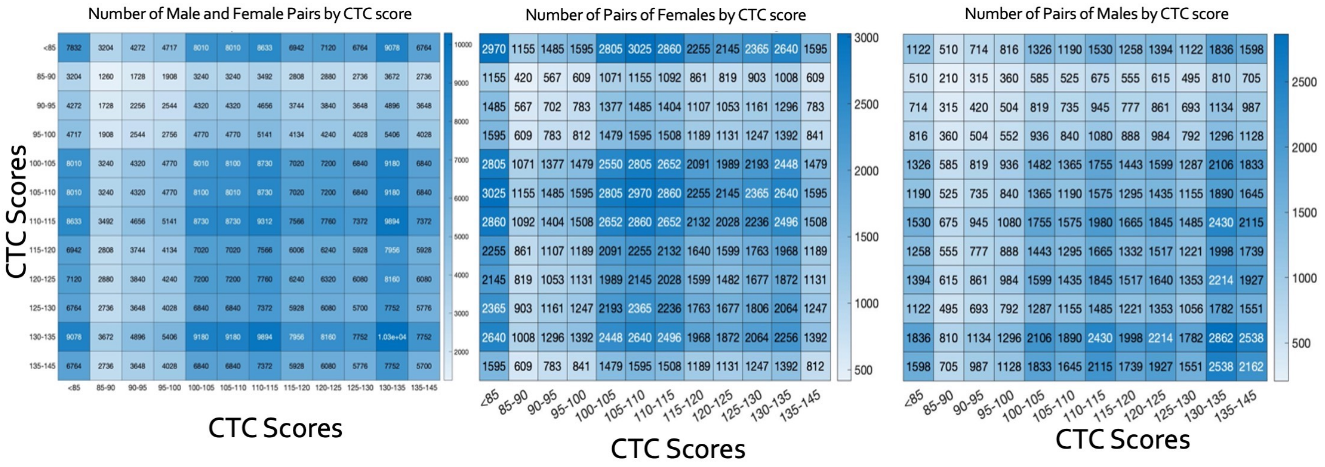


**Figure S6.** Number of pairs of individuals for each combination of CTC scores for all pairs (left), female-female pairs (middle) and male-male pairs (right).
